# An Exact and Fast SAT Formulation for the DCJ Distance

**DOI:** 10.1101/2024.11.05.622153

**Authors:** Aaryan M. Sarnaik, Ke Chen, Austin Diaz, Mingfu Shao

**Affiliations:** Department of Computer Science and Engineering, The Pennsylvania State University, University Park, PA 16802, USA; Huck Institutes of the Life Sciences, The Pennsylvania State University, University Park, PA 16802, USA

**Keywords:** SAT, ILP, DCJ, MCD

## Abstract

Reducing into a satisfiability (SAT) formulation has been proven effective in solving certain NP-hard problems. In this work, we extend this research by presenting a novel SAT formulation for computing the double-cut-and-join (DCJ) distance between two genomes with duplicate genes. The DCJ distance serves as a crucial metric in studying genome rearrangement. Previous work achieved an exact solution by transforming it into a maximum cycle decomposition (MCD) problem on the corresponding adjacency graph of two genomes, followed by reducing this problem into an integer linear programming (ILP) formulation. Using both simulated datasets and real genomic datasets, we firmly conclude that the SAT-based method scales much better and runs faster than using ILP, being able to solve a whole new range of instances which the ILP-based method cannot solve in a reasonable amount of time. This underscores the SAT-based approach as a versatile and more powerful alternative to ILP, with promising implications for broader applications in computational biology.

## 1 Introduction

Genome rearrangements are large-scale evolutionary events that change the structural organization of a genome. It is known that genome rearrangements are associated with various diseases, including cancers, congenital disorders, and neurodevelopmental conditions [8]. Studying these rearrangements may identify specific genetic changes that contribute to diseases, aiding in diagnostics and targeted therapies. They also act as a metric to measure evolutionary distance between species [19]. Genomic rearrangements include inversions, transpositions, circularizations, and linearizations, all of which act on a single chromosome, and translocations, fusions, and fissions, which act on two chromosomes. While individual rearrangement events are of interest for certain scenarios, mathematical models have been proposed to unify multiple arrangement events [6]. Among these, the most successful yet simple model is arguably the double-cut-and-join (DCJ) operation [4,31]. A DCJ operation makes two cuts in the genome, either in the same chromosome or in two different chromosomes, producing four cut ends, then rejoins the four cut ends. Each of the genomic rearrangements mentioned above can be characterized by a single DCJ operation.

The advent of whole-genome sequencing has provided us with masses of data on which to study genomic rearrangements. A fundamental computational problem involved is to compute the edit distance between two genomes, i.e., the minimum number of evolutionary events needed to transform one genome into another. Rich algorithmic research has been conducted in this direction with also fruitful applications [5,9,10,32,21]. Under the inversion model, Hannenhalli and Pevzner pioneered the first polynomial-time algorithm to compute the edit distance for unichromosomal genomes [15], which was later improved to linear time [3] and extended to multichromosomal genomes [15,18,24,29,5]. Under the DCJ model, the edit distance can be computed in linear time in a very simple and elegant way [4]. However, when genomes contain duplicated genes, which is the case for most genomes, most above edit distance problems become NP-hard [11]. Efficient heuristics such as MSOAR [28,13] have been designed to solve the inversion distance. Exact algorithms for various edit distance have also been designed [11]. Most of the exact algorithms transformed the edit distance problem into integer linear programming (ILP), followed by calling existing ILP solvers [17,20]. Examples include the first ILP formulation for the DCJ distance [27], and ILP formulations for various edit distance problems [7].

These ILP formulations cannot scale. It is easy for the solver to reach a pre-determined time limit (for example, a day). More scalable, exact algorithms are therefore desirable. A promising direction is the reduction into a satisfiability (SAT) problem followed by calling a SAT solver [14]. The SAT problem is one of the most fundamental problems in computer science. A SAT instance consists of a set of binary variables and a set of clauses, each of which is a disjunction (i.e., or relationship) of variables or their negations, and the problem seeks whether there exists an assignment to variables so that all clauses can be satisfied. SAT problems (for example 3-SAT) are proved to be NP-complete, granting it general power to solve other problems like ILP. Moreover, SAT problems are light-weight, with excellent solvers available that can solve larger instances swiftly [16]. SAT solvers have demonstrated particularly impressive performance characteristics compared to ILP approaches, especially for highly combinatorial problems like genome rearrangements. While ILP solvers must manage continuous relaxations and branch-and-bound trees, SAT solvers can focus purely on boolean constraint propagation and conflict-driven clause learning, often leading to improvement in orders of magnitude in solving time for suitable problems. This efficiency gain becomes particularly crucial when analyzing complex genomic datasets with numerous duplicated genes. There are already successful applications of SAT solvers in computational biology [14], demonstrating the advantages of transforming problems into SAT problems.

In this work, we explore if the DCJ distance for genomes with duplicate genes can be more efficiently solved by reducing it into a SAT problem. To do this, we derived a novel SAT formulation on the adjacency graph of the two genomes that is equivalent to finding the DCJ distance between them. This is non-trivial given the need of counting the number of cycles with variables. We compare the runtime of solving the state-of-the-art ILP formulation and our SAT formulation, using both simulated and biological datasets. The results demonstrate that our SAT formulation scales much better, being able to solve a whole new range of complex problems in time that is several orders of magnitude shorter than ILP. This not only pushed forward the boundaries of solvable DCJ instances but also showcased another successful example of using SAT solving methods, shedding light on its use for broader applications in bioinformatics.

## 2 Preliminaries

In studying genomic rearrangements, a genome is represented as a set of chromosomes and each chromosome is represented as a linear or circular list of genes. An example is *G* = ⟨*a*^1^, *b*, −*a*^2^⟩(−*c, d*), which contains a linear chromosome enclosed by ⟨⟩ and a circular chromosome enclosed by (). Each gene *g* can be thought of an arrow, pointing from its tail *g*_*t*_ to its head *g*_*h*_, which together are referred to as its *extremities*. The sign of a gene represents its transcriptional direction; we usually pick the left to right direction as positive, then a negative sign indicates the head of the gene is to the left of its tail. Two consecutive genes in a chromosome can be represented as an *adjacency* which consists of two adjacent extremities, one from each of the genes. The ends of a linear chromosome are called *telomeres*, consisting of a single extremity. For example, the linear chromosome in *G* contains two telomeres 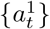 and 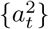, and two adjacencies 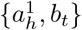 and 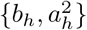. Homologous genes are grouped into gene family; they are labeled with the same letter but distinguished with different superscripts. For homologous genes *a*^*i*^ and *a*^*j*^, their corresponding extremities (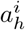 and 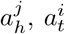 and 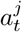) are also called homologous.

Given two genomes *G*_1_ and *G*_2_ with the same gene content and without duplicate genes (i.e., each gene family has exactly one copy in each genome, the DCJ distance between them, denoted as *d*(*G*_1_, *G*_2_), can be simply calculated using the *adjacency graph A* = (*V*_1_ ∪ *V*_2_, *E*_*g*_ ∪ *E*_*b*_). *V*_1_ consists of all *extremities* in *G*_1_, and *V*_2_ consists of all extremities in *G*_2_. The two extremities in one adjacency are connected with a *gray edge* in *E*_*g*_, and homologous extremities across the two genomes are connected with *black* edge in *E*_*b*_. See Fig. 1 (Left) for an example. When *G*_1_ and *G*_2_ do not contain duplicate genes, each vertex has degree one or two, hence the adjacency graph consists of vertex-disjoint paths and cycles. The DCJ distance can be simply calculated as *d*(*G*_1_, *G*_2_) = |*V*_1_ ∪ *V*_2_|*/*4 − *c* − *p/*2, where *c* is the number of cycles in *G*, and *p* is the number of paths in *A* [4].

**Fig. 1:**
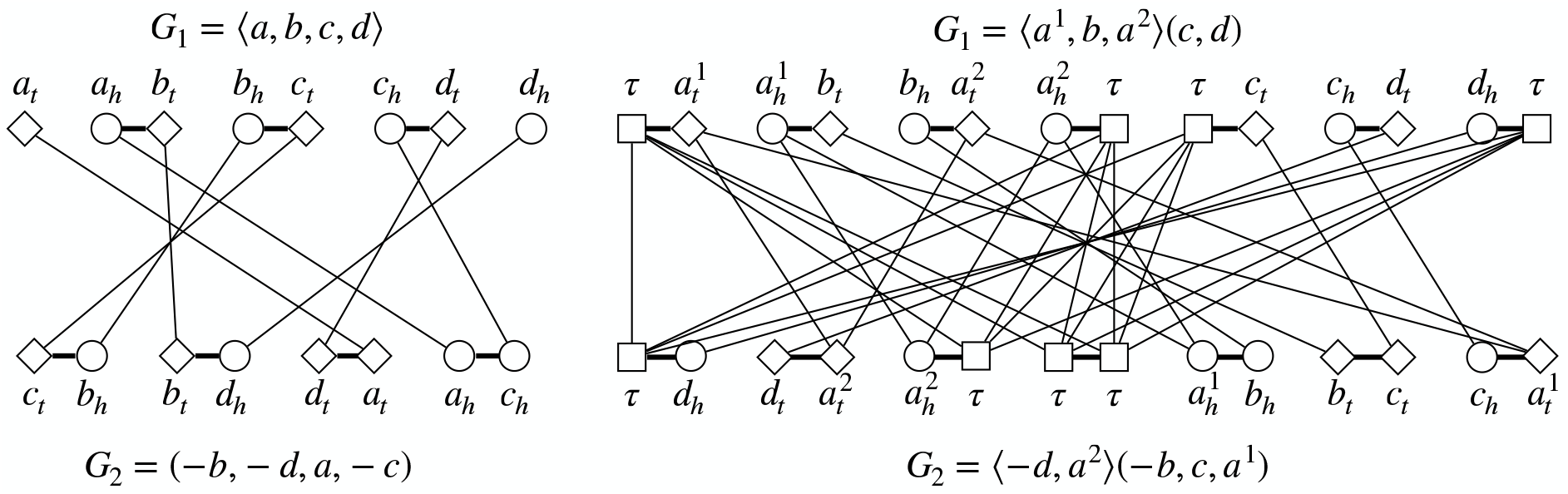
Left: The adjacency graph between two genomes without duplicate genes. Right: The adjacency graph for two genomes with duplicate genes and null extremities. In each graph, the grey edges are depicted by bold horizontal bars; the black edges run between the top and bottom genomes. The added null extremities *τ* connect to all null extremities on the opposite side.

Now consider two genomes *G*_1_ and *G*_2_ with the same gene content but containing duplicate genes. To construct the corresponding adjacency graph *A* = (*V*_1_ ∪ *V*_2_, *E*_*g*_ ∪ *E*_*g*_), we add a *null extremity*, denoted as *τ*, to each telomere to make it an adjacency. If the number of null extremities added to the two genomes are not balanced, we will add adjacencies {*τ, τ* } to make it balanced. After that, the construction of adjacency graph *A* is the same as that for genomes without duplicate genes. See Fig. 1 (Right) for an example. Note that the extremities of the duplicate genes have degrees more than two (they connect to each homologous extremities in the opposite genome), the adjacency graph can no longer be covered by a set of vertex-disjoint paths and cycles.

The problem of computing the DCJ distance between two genomes with duplicate genes can be reduced to the problem of finding the optimal consistent decomposition of the adjacency graph [27]. A *decomposition* of the adjacency graph *A* is a set of vertex-disjoint alternating cycles that cover all vertices in *A*, where *alternating cycle* means any two adjacent edges in it consist of one black edge and one gray edge. Note that because of the alternating requirement, covering all the vertices is equivalent to covering all the gray edges in the decomposition. A decomposition is *consistent* if for any two homologous genes *a*^*i*^ and *a*^*j*^, either both black edges 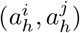 and 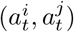 are in the decomposition or neither of them are in the decomposition. Given a consistent decomposition with *c* cycles, exactly (|*V*_1_ ∪ *V*_2_|*/*4 − *c*) DCJ operations are needed to transform *G*_1_ into *G*_2_. Hence, minimizing the number of DCJ operations needed is equivalent to finding a consistent decomposition in *A* with maximized total number of cycles. Formally, *d*(*G*_1_, *G*_2_) = min_*D*∈𝒟_(|*V*_1_∪*V*_2_|*/*4−*c*_*D*_) = |*V*_1_ ∪ *V*_2_|*/*4 −max_*D*∈𝒟_(*c*_*D*_), where 𝒟 is the set of all consistent decompositions and *c*_*D*_ represents the number of cycles in *D*.

## 3 SAT Formulation

We transform the above maximum-cycle decomposition problem into a SAT problem. The resulting SAT formulation is in conjunctive normal form (CNF), consisting of a conjunction of clauses where each clause is a disjunction of literals (boolean variables or their negations). There is no requirement on the number of literals in each clause.

### ALO and AMO

We first introduce a technique that will be used multiple times in the following derivation. Let *x*_1_, *x*_2_, · · ·, *x*_*n*_ be *n* literals. We often need to enforce that at least one (ALO) of them must be TRUE, denoted as *ALO* (*x*_1_, *x*_2_, · · ·, *x*_*n*_), or require that at most one (AMO) of them is TRUE, denoted as *AMO* (*x*_1_, *x*_2_, · · ·, *x*_*n*_). ALO can be easily implemented using one clause in CNF form:

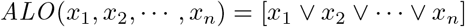

A straightforward way to implement AMO is to use pairwise encoding [22,23], which can be implemented with *O*(*n*^2^) clauses: (¬*x*_*i*_ ∨ ¬*x*_*j*_), for all 1 ≤ *i* < *j* ≤ *n*. To reduce the number of clauses, we adapt a technique called binary encoding proposed by [12], which has been used to successfully solve a number of large SAT instances with a standard SAT local search method [26]. This encoding uses *O*(*n* log *n*) clauses with additional *O*(log *n*) boolean variables: *y*_1_, *y*_2_, · · ·, *y*_*k*_, where *k* = ⌈log *n*⌉.AMO can be implemented as:

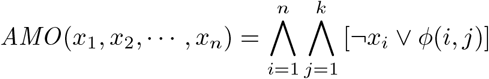

where *ϕ*(*i, j*) = *y*_*j*_ if the *j*-th bit of binary representation of number *i* − 1 is 1, and *ϕ*(*i, j*) = ¬*y*_*j*_ otherwise. The idea is to create different sequences of ⌈log *n*⌉-tuples *y*_*j*_, 1 ≤ *j* ≤ ⌈log *n*⌉, such that whenever *x*_*i*_ is assigned TRUE, 1 ≤ *i* ≤ *n*, then we can infer that the other variables *x*_*i*_*′* must be FALSE, for any *i′*≠*i*.

### Ensuring a Decomposition

We introduce boolean variables *x*_*e*_ for each edge *e* ∈ (*E*_*g*_ ∪ *E*_*b*_), representing whether that edge will be chosen in the final decomposition *D*: *x*_*e*_ is TRUE means *e* is selected. To ensure that *D* includes all the gray edges, for each gray edge *e* ∈ *E*_*g*_, we add the single-literal clause ([*x*_*e*_]) to the formulation. Adding a single-literal clause forces the literal to be TRUE. For each vertex *v* ∈ *V*_1_ ∪ *V*_2_, we also need to ensure that exactly one black edge adjacent to *v* is selected. This is equivalent to selecting at least one AND at most one adjacent black edge to *v*, which therefore can be implemented by adding the following clauses:

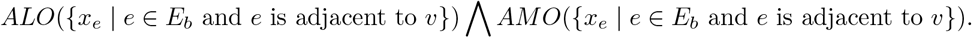

### Ensuring A Consistent Decomposition

For every pair of homologous genes (*a*^*i*^, *a*^*j*^) across the two genomes, we must ensure that black edge 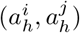 is chosen if and only if its homologous counterpart 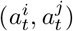 is chosen, i.e., ensuring 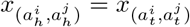. We translate it into CNF form using simple logical rules and add the simplified CNF formula as clauses:

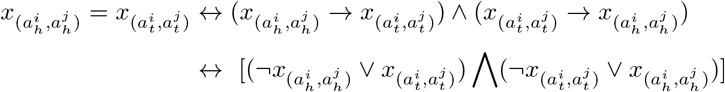

### Counting Cycles

The above variables and clauses encode and enforce a consistent decomposition. The key challenge is to count the number of cycles in it and maximize it. We follow an idea introduced in [27] as part of the ILP formulation, that uses “labels” to count cycles. Note that integer variables in ILP are naturally suited for counting, we must overcome extra challenges to accomplish the same in the SAT formulation.

We rename the vertices as *V*_1_ = {*v*_1_, *v*_2_, …, *v*_*n*_} and *V*_2_ = {*v*_*n*+1_, *v*_*n*+2_, …, *v*_2*n*_}. For each vertex *v*_*i*_ ∈ *V*_1_ ∪ *V*_2_, we introduce *i* new boolean variables {*w*(*i, t*) | 1 ≤ *t* ≤ *i*} to assign *v*_*i*_ an integral label in the range {1, 2, · · ·, *i*}. If some *w*(*i, t*) is TRUE, it means that vertex *v*_*i*_ is assigned the label *t*, indicating it is covered by the *t*-th cycle. We first ensure that, for each *v*_*i*_ ∈ *V*_1_ ∪ *V*_2_, exactly one variable in {*w*(*i, t*)} is TRUE, which can be done by adding the set of clauses:

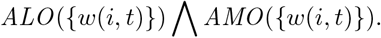

We then ensure that all vertices in the same cycle of the resulting decomposition *D* share the same labeled. This can be guaranteed by ensuring that, if an edge is chosen, then its two incident vertices must have the same label. Formally, this is equivalent to saying that, if 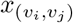 is TRUE, then *w*(*i, t*) = *w*(*j, t*) for all 1 ≤ *t* ≤ min{*i, j*}. One can verify that 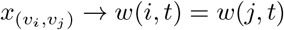 can be implemented in CNF form as:

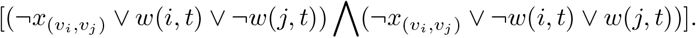

We therefore add the above two clauses to the SAT formulation for every edge (*v*_*i*_, *v*_*j*_) ∈ *E*_*g*_ ∪ *E*_*b*_ and for every *t*, 1 ≤ *t* ≤ min{*i, j*}.

Note that the upper bound of the label for *v*_*i*_ is *i*, because we can always label a cycle by the smallest-indexed vertex it covers so that the label of each vertex (i.e., the cycle it is in) is at most its index. Furthermore, by this labeling, the number of vertices that have the same index and label is exactly the number of cycles, which is what we aim to count. We introduce a boolean variable *z*_*i*_ for each vertex *v*_*i*_ ∈ *V*_1_ ∪ *V*_2_, 1 ≤ *i* ≤ 2*n*, to indicate if the label of *v*_*i*_ is equal to its index *i*. Formally, *z*_*i*_ is TRUE only if *w*(*i, i*) is TRUE, which can be implemented with one clause:

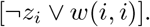

We adapt an approach described in [14] to count the number of *z*_*i*_’s that are TRUE. We introduce boolean variables *T* (*i, d*), for 1 ≤ *i* ≤ 2*n* + 1 and 0 ≤ *d* ≤ *i* − 1, that is TRUE if and only if at least *d* variables in the set {*z*_*j*_ | *j* ≤ *i* − 1} are TRUE. We first enforce this definition by adding clauses in an inductive way. For the base cases, we add single-variable clause:

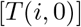

for all 1 ≤ *i* ≤ 2*n* + 1, to ensure that at least 0 variables in {*z*_*j*_ | *j* ≤ *i* − 1} are TRUE. We then enforce that, if *T* (*i, d*) is TRUE then *T* (*i* + 1, *d*) must be TRUE, which can be implemented as one clause:

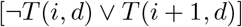

and we add this clause to the SAT formulation for all 1 ≤ *i* ≤ 2*n* + 1 and 0 ≤ *d* ≤ *i* − 1.

Next, we enforce that if *T* (*i, d*) is TRUE AND *z*_*i*_ is TRUE, then *T* (*i* + 1, *d* + 1) must be TRUE. This can be implemented by the clause:

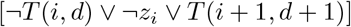

and again we add this clause for all 1 ≤ *i* ≤ 2*n* + 1 and 0 ≤ *d* ≤ *i* − 1.

Last, we need to ensure the “only if” side of the logic, i.e., if *T* (*i*+ 1, *d*) is TRUE, then either *T* (*i, d*) is TRUE, or both *T* (*i, d* − 1) and *z*_*i*_ are TRUE. Formally, this implies *T* (*i* + 1, *d*) → [*T* (*i, d*) ∨ (*T* (*i, d* − 1) ∧ *z*_*i*_)]. Trying to simplify this formula to CNF form using simple rules would lead to an exponential number of clauses. Hence, we use Tseytin encoding [25,30] to convert it to CNF form with fewer clauses. We introduce auxiliary boolean variables *Q*(*i, d*), for all 1 ≤ *i* ≤ 2*n* + 1 and 1 ≤ *d* ≤ *i* + 1, defined as *Q*(*i, d*) = *T* (*i, d* − 1) ∧ *z*_*i*_. This definition can be implemented in CNF form as:

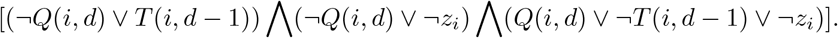

The remaining logic *T* (*i* + 1, *d*) → *T* (*i, d*) ∨ *Q*(*i, d*) can be simply implemented as:

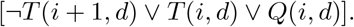

### Maximizing Number of Cycles

Our SAT formulation so far has the ability to make consistent decompositions and also to count the number of cycles in the result. The last step is to add a single-variable clause

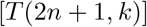

as a “query”. If the resulting SAT instance is satisfiable, then it means a consistent decomposition with *k* cycles exists but the optimal solution may contain more cycles. To find the maximum-cycle decomposition, we add the clauses [*T* (2*n* + 1, *k*)] to the formulation one by one starting from *k* = 1, until an unsatisfiable instance is reached at *k* = *k*^*∗*^. At this point, we can conclude that the maximum number of cycles in a consistent decomposition is *k*^*∗*^ − 1.

Note that a seemingly better choice here is to apply binary search instead of the linear scan as described above. However, we observe that 1. the unsatisfiable instance is usually the most time consuming one to solve; and 2. modern SAT solvers support “warm start” that makes solving consecutive instances with small changes (namely, from *k* to *k* + 1) rather efficient. The following experimental results demonstrate that this linear scan is indeed a feasible and effective strategy.

## 4 Results

### Pipeline

We compare the computational efficiency of our SAT formulation against the state-of-the-art ILP formulation. The experiments are carried out on both simulated instances (Section 4.1) and real genomes (Section 4.2). For an instance, which consists of a pair of genomes *G*_1_ and *G*_2_, the corresponding adjacency graph *A* is constructed first (Section 2). We then apply three pre-processing techniques to simplify *A* without affecting optimality. The simplified graph, denoted as *A′*, is then transformed into the SAT formulation, described in this paper, or to the ILP formulation, described in [27] and implemented in the GREDU package. The ILP formulation is solved using the GUROBI solver [17]. Our SAT formulation is solved using the Glucose4 solver [2] in the PySAT library [16]. All tests are conducted using a single thread on an Intel(R) Xeon(R) Gold 6230 CPU @ 2.10GHz. We measure the time of each solver from being given the simplified graph to returning the optimal number of cycles. We perform all experiments with a time limit of 9 hours (43200 seconds); the actual time (user time + system time) is recorded if a method finishes within the time limit; otherwise, a TLE (time limit exceeded) is reported.

### Pre-processing

There are some pre-processing techniques that can be used to reduce the complexity of the graph while preserving optimality [27]. We simplify the adjacency graph using these techniques and then give the processed graph as input to both the SAT and ILP formulations. Below, the length of a cycle is defined to be the number of black edges in it; the degree of a vertex is the total number of gray edges and black edges it is incident to.

1. A cycle of length two can be “fixed” if the cycle contains some vertex with degree 2, see Fig. 2 (Left). To fix a cycle, for each vertex in it, we keep the incident black edges that are part of the cycle and remove other (if any) incident black edges. For any two homologous genes *a*^*i*^ and *a*^*j*^ across two genomes, if 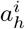 and 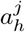 gets fixed in a length-two cycle, then its other pair 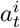 and 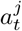 are also fixed.
2. An alternating path that starts and ends at two vertices with a degree of 2 (say, *u* and *v*) and only contains vertices with degree 2 can be shrunk down to a single black edge between *u* and *v*, provided that *u* and *v* are connected by a gray edge to a vertex with degree > 2 and *u* and *v* lie in different genomes. See Fig. 2 (Middle).
3. An alternating path that starts and ends at two vertices with a degree > 2 (say, *u* and *v*) and only contains vertices with degree 2 (except *u* and *v*) can be shrunk down to a single gray edge between *u* and *v*, provided that *u* and *v* lie in the same genome. See Fig. 2 (Right).

**Fig. 2:**
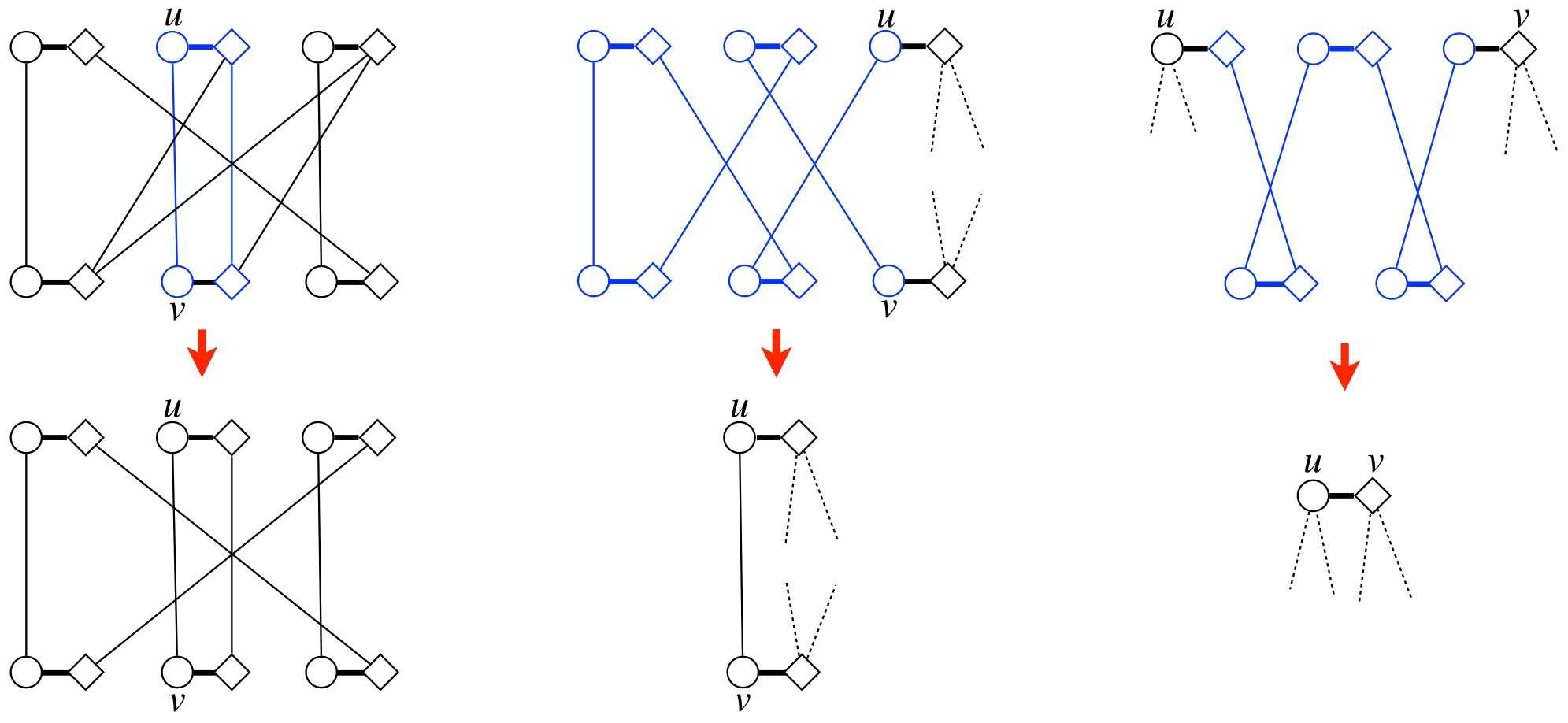
Left: simplifying using a length-two cycle, in which *u* and *v* have degree 2. Middle and Right: shrinking alternating paths.

### 4.1 Results on Simulated Data

We first evaluate the ILP formulation and the SAT formulation on simulated data. The simulation is governed by 3 parameters, the number of genes *N*_*g*_, the number of gene families *N*_*f*_, and the number of DCJ operations *N*_*d*_. We start with simulating a random ancestral genome with *N*_*g*_ genes and *N*_*f*_ gene families by specifying the number of genes, number of chromosomes, and number of gene families. To simulate a pair of derived genomes, we independently perform *N*_*d*_ random DCJ operations (in terms of reversals) on the ancestral genome twice. The complexity of the resulting adjacency graph, and consequently, the complexity of the maximum cycle decomposition problem, varies as the three parameters vary. For a fixed number of genes *N*_*g*_, as the number of gene families *N*_*f*_ decreases, the number of duplicate genes increases, thereby increasing the complexity. Also, as the number of DCJ operations *N*_*d*_ increases, the pre-processing will be less effective in simplifying the adjacency, leading to more complex instances to solve.

We perform two sets of experiments with *N*_*g*_ ∈ {100, 500, 1500, 5000}. In the first experiment, we keep the number of gene families *N*_*f*_ fixed and gradually increase the number of DCJ operations *N*_*d*_ performed. In the second experiment, we vary the number of gene families *N*_*f*_. The results for these two experiments are illustrated in Fig. 3 and Fig. 4, respectively.

**Fig. 3:**
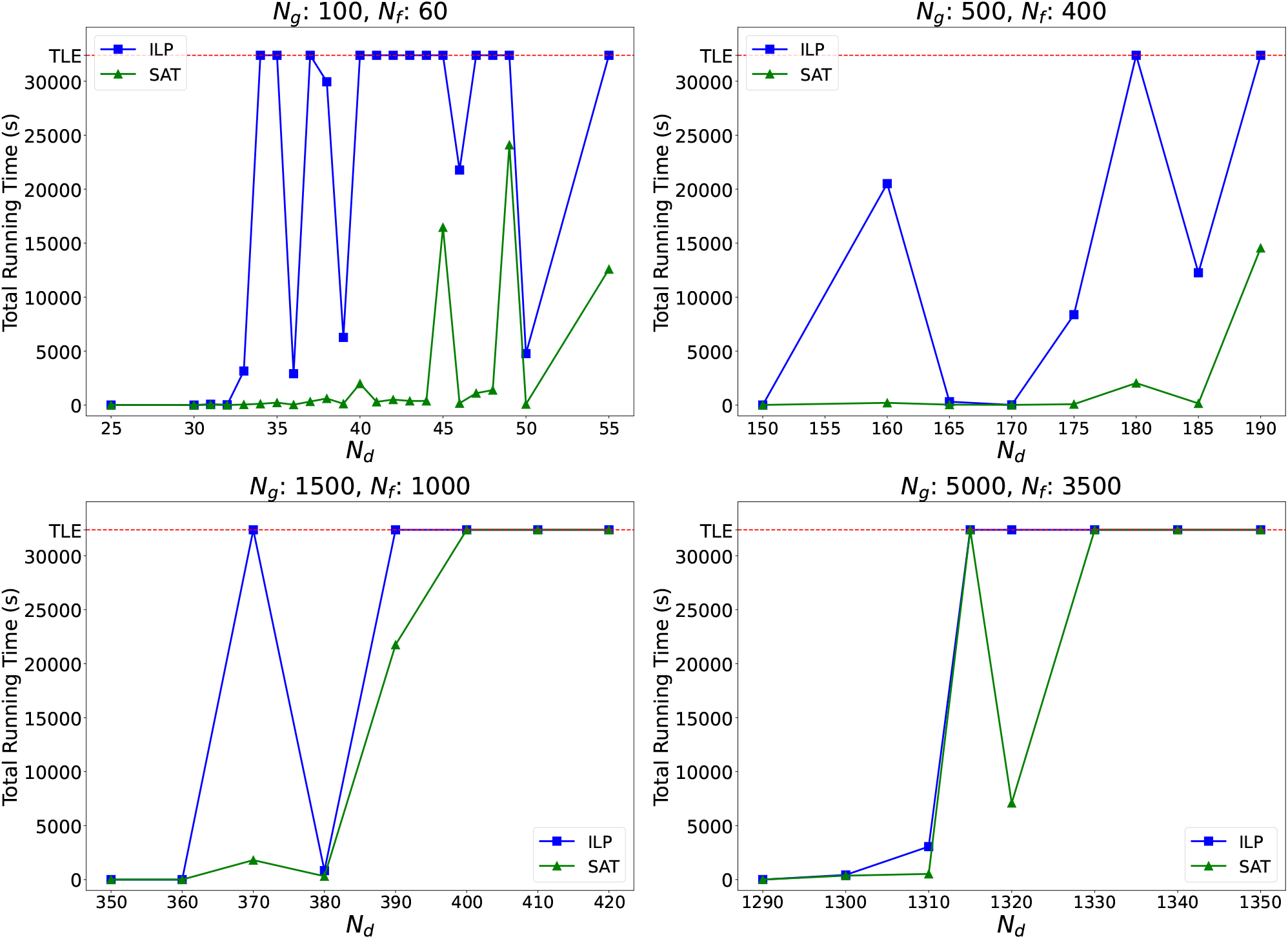
Comparison of runtime (in seconds) of ILP and SAT formulations with simulated data. Each panel corresponds to a fixed *N*_*g*_ and *N*_*f*_ with varying number of DCJ operations. Each point on the graph represents an instance. The ILP formulation is represented by the blue squares and the SAT formulation is represented by the green triangles.

**Fig. 4:**
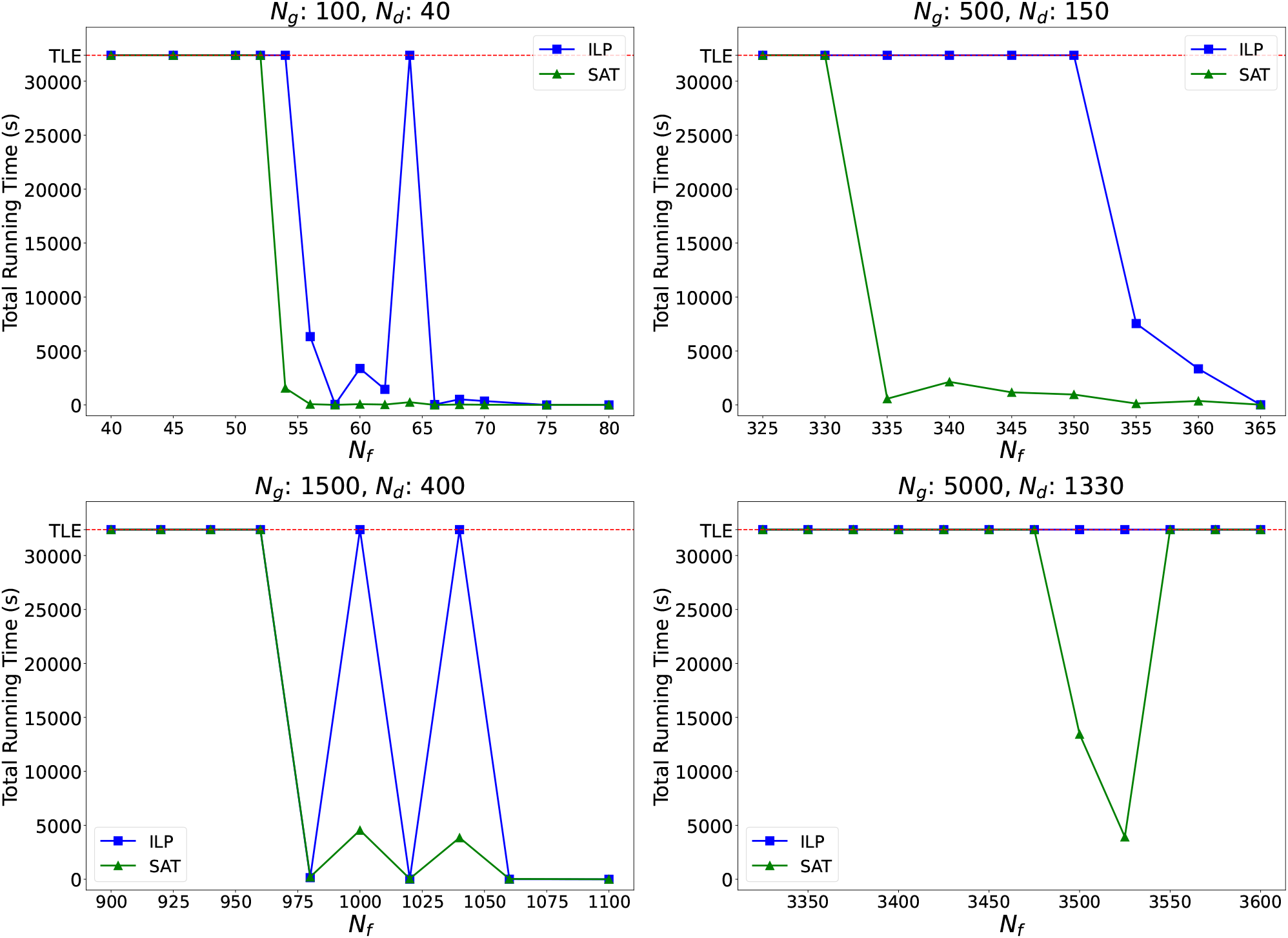
Comparison of runtime (in seconds) of ILP and SAT formulations with simulated data. Each panel corresponds to a fixed *N*_*g*_ and *N*_*d*_ with varying number of gene families. Each point on the graph represents an instance. The ILP formulation is represented by the blue squares and the SAT formulation is represented by the green triangles.

The first and the foremost observation is that, the SAT formulation is consistently more efficient than the ILP formulation on all instances that are solvable within the time limit. This firmly demonstrates the superiority of using SAT instead of ILP. Importantly, over the total 93 instances, SAT can solve 66 of them within the time limit whereas ILP solves only 38 of them within the time limit. This shows that by using SAT formulations the range of tractable instances can be substantially expanded.

The second observation (not only for simulated data, but also for real data) is that, in each of the simulation settings the instances can be roughly divided into three sections on the basis of the complexity of the instances. The first section consists of the easily solvable instances which both methods can finish in at most a few seconds. The second section consists of a range of more complex graphs, where SAT exceedingly outperforms ILP. The third section consists of extremely complex problems which are intractable for both methods, leading to an inflection point at which both methods exceed the time limits. This is an expected result from solving an NP-Hard problem: once the complexity of instances increases to a certain point, no algorithm can find exact solution in an acceptable amount of time. The existence of the middle section shows the striking ground that SAT covers, which ILP can’t, pushing forward the boundaries of tractable instances.

Specifically, in Fig. 3, the power of the SAT formulation is most apparent in the genomes with 100 genes and 60 gene families, and 500 genes and 400 gene families. For lesser number of DCJ operations, both methods are able to find the optimal cycles very fast, hence we observe that the corresponding points are very close together and have total time close to 0. However, as soon as we increase the DCJ operations, ILP starts failing to solve them (becomes TLE) or solves them in a time much more than the time taken by using SAT. For these smaller genomes, we do not see clearly an inflection point (but the overall trend is there). For the genomes with 1500 genes and 5000 genes, we are able to distinguish the three sections described earlier clearly. Some bounces (solvable → TLE → solvable, for example, the bouncing of ILP on 1500 genes and SAT on 5000 genes) are observed which can be accounted for by the random approach we used when performing these DCJ operations in the larger graphs. We also notice this trend in Fig. 4. As the number of gene families decreases in the genomes, the problem goes from (solvable by both) to (solvable only by SAT) to (infeasible). We still have instances that do not exactly adhere to this trend, but these can again be accounted by the randomness of the generation of the genomes.

### 4.2 Results on Real Data

We selected a set of 7 well-annotated species (human, gorilla, mouse, macaque, chicken, zebrafish, molerat) and for each species, we downloaded their annotated gene data for all protein-coding genes from Ensembl [1], including gene family names, positions on the chromosomes and gene symbols. If a gene has multiple alternative products, we keep its longest isoform. Two genes are considered homologous if they have the same Ensembl gene family name. Two genes are considered orthologous if they have the same gene symbol. Note that two orthologous genes are necessarily homologous, but two homologous genes need not be orthologous. Each gene in one genome will have at most one orthologous gene in another genome.

We perform pairwise comparison over the 7 genomes. For each pair, we derive a pair of genomes with the same gene content by keeping only orthologous gene pairs. However, the resulting adjacency graphs for each pair remains fairly large, which turns out to be intractable for both methods (apart from 2 pairs which can be solved by SAT; see “All” column of Table 1). To get a better measurement, we further simplify these instances by filtering out the gene families that have a count in each genome greater than 2 and 3 (see columns “≤ 2” and “≤ 3” in Table 1 respectively). Note that by doing so most genes can still be kept. This leads to simplified instances and consequently, comparatively simplified graphs and allows us to better analyze the results.

**Table 1:** Comparison of runtime of SAT and ILP over 21 pairs of genomes. Bold numbers highlight instances where formulation can solve but not ILP. Italic numbers highlight instances where ILP runs faster than SAT. The pairs are sorted according to the runtime of SAT in the “≤ 2” column.

| Genome Pair | $\leq 2$ | | $\leq 3$ | | All | |
| --- | --- | --- | --- | --- | --- | --- |
|  | ILP | SAT | ILP | SAT | ILP | SAT |
| Gorilla vs. Human | 0.351 | 0.160 | 0.494 | 0.341 | **TLE** | **2571.492** |
| Human vs. Macaque | 0.393 | 0.275 | *0.609* | *0.886* | TLE | TLE |
| Macaque vs. Mouse | 0.609 | 0.348 | *1.077* | *1.634* | **TLE** | **14771.563** |
| Gorilla vs. Macaque | *0.399* | *0.552* | 26.210 | 16.503 | TLE | TLE |
| Human vs. Mouse | 2.069 | 1.149 | 2.489 | 2.298 | TLE | TLE |
| Molerat vs. Mouse | 5.427 | 1.800 | **TLE** | **29.880** | TLE | TLE |
| Macaque vs. Molerat | 18.721 | 1.949 | **TLE** | **2283.746** | TLE | TLE |
| Human vs. Molerat | **TLE** | **2.548** | **TLE** | **9.518** | TLE | TLE |
| Gorilla vs. Molerat | **TLE** | **4.097** | TLE | TLE | TLE | TLE |
| Gorilla vs. Mouse | 7194.601 | 9.723 | **TLE** | **34.254** | TLE | TLE |
| Chicken vs. Macaque | **TLE** | **12.973** | **TLE** | **1509.596** | TLE | TLE |
| Chicken vs. Human | 25.715 | 13.720 | **TLE** | **242.045** | TLE | TLE |
| Chicken vs. Mouse | **TLE** | **50.232** | **TLE** | **20746.844** | TLE | TLE |
| Chicken vs. Gorilla | **TLE** | **152.967** | TLE | TLE | TLE | TLE |
| Chicken vs. Molerat | **TLE** | **401.713** | TLE | TLE | TLE | TLE |
| Macaque vs. Zebrafish | TLE | TLE | TLE | TLE | TLE | TLE |
| Human vs. Zebrafish | TLE | TLE | TLE | TLE | TLE | TLE |
| Mouse vs. Zebrafish | TLE | TLE | TLE | TLE | TLE | TLE |
| Gorilla vs. Zebrafish | TLE | TLE | TLE | TLE | TLE | TLE |
| Molerat vs. Zebrafish | TLE | TLE | TLE | TLE | TLE | TLE |
| Chicken vs. Zebrafish | TLE | TLE | TLE | TLE | TLE | TLE |

The detailed comparison is given in Table 1. For “≤ 2”, “≤ 3”, and “All” instances, there are 6, 7, and 2 instances, respectively, where SAT can solve but ILP cannot (highlighted in **bold**). This again proves that using SAT formulations scales better than using ILP formulations. For those instances (14 of them) that both methods can solve, SAT also runs much faster (except 3 pairs where both methods can finish in less than 2 seconds). These results align very well with the results we show using simulations.

## 5 Conclusion and Discussion

Our work demonstrates that SAT-based approaches can significantly advance the computational capabilities in solving the DCJ distance problem, a fundamental challenge in comparative genomics. The experimental results clearly show that our SAT-based formulation not only maintains the exactness of previous ILP-based methods but substantially expands the feasible problem space, successfully handling instances that were previously computationally intractable. This improvement is particularly evident in the “middle ground” of problem complexity, where SAT consistently outperforms ILP approaches, solving problems that would otherwise timeout under ILP formulation. The success of this SAT-based approach has broader implications beyond the specific domain of DCJ distance calculation. It suggests that other NP-hard problems in computational biology, particularly those currently addressed through ILP formulations, might benefit from similar SAT-based reformulations. This could potentially unlock new avenues of research by making previously unfeasible analyses possible, especially when dealing with large-scale genomic data.

Deriving the SAT formulation can be complex. In this work, we integrated several techniques, including AMO constraints, cycle counting using labeling, Dan Gusfield’s method [14], and Tseytin encoding, to develop a functional formulation. Future work will also focus on reducing the number of clauses and variables or finding a better balance between them, to achieve a more efficient formulation leading to even better results and to scale to even larger instances.

## Availability

The source code and the scripts that can reproduce the experimental results in this paper is available at https://github.com/Shao-Group/dcj-sat.

## Acknowledgment

This work is supported by the US National Science Foundation (2145171 to M.S.) and by the US National Institutes of Health (R01HG011065 to M.S.).

